# A reservoir model of respiration-induced perceptual alternation in binocular rivalry

**DOI:** 10.1101/2025.02.06.636993

**Authors:** Shogo Yonekura, Atsushi Narita, Hoshinori Kanazawa, Yasuo Kuniyoshi

## Abstract

Perceptual alternation in human binocular rivalry is more likely to occur during certain respiratory phases. In this paper, we show that the respiration dependence of perceptual alternations can be reproduced by a randomly connected recurrent neural network coupled with respiration relevant information via a neuromodulator of norepinephrine (NA). We considered two models of NA modulations; NA increases or decreases the nonlinearity of the activation function of neurons, and we found that the shape of the likelihood function of perceptual alternation depends only on respiratory phase, regardless of whether NA increases or decreases neural nonlinearity. Our results suggest that periodic neuromodulation facilitates the switching of competing neural states in specific phases and that this effect is independent of the excitatory or inhibitory effect of NA.

## 1 Introduction

The extensive range of brain activity and cognitive processes are coordinated with the physiological processes occurring within the body’s internal organs, including respiration, gastric peristalsis, and cardiac rhythm [8, 9, 17, 25, 26, 29, 32, 38]. Recent evidence has additionally indicated that perceptual alternation in human binocular rivalry is dependent on respiration [36].

Moreover, the recent advancements in computational neuroscience have demonstrated that a randomly connected recurrent neural network, i.e., a reservoir network, is capable of approximating an arbitrary function through the use of a linear readout function, despite the random and fixed internal synaptic connections [12, 13, 22]. This new computational scheme, which does not require updating the internal synaptic weight, is called reservoir computing. It is further assumed that the local cortical microcircuit as well as the global cortical network can be viewed as reservoir networks [16, 21, 31]. It is well documented that binocular rivalry is the result of the interaction of numerous brain regions, including the lateral geniculate nucleus, the thalamic reticular nucleus, and the visual areas V1, V2, V4, MT, and IT. [14].

Therefore, it may be reasonable to assume that respiratory-synchronized perceptual alternation is the product of the interaction among the global reservoir networks and respiratory activity.

The present study demonstrates that the dependency of perceptual alternation on respiratory phase can be generated by a reservoir network, wherein the nonlinearity of the activation function of neurons is modulated based on respiratory dynamics.

## 2 Model

As we discussed above, binocular rivalry is generated by the interaction of many brain regions [14]. Moreover, recent developments in computational neuroscience allow us to consider the global neural network as a large-scale reservoir network [31]. Therefore, we use an echo state network (ESN) [12, 13] as a rough model of a global recurrent neural network that correlates with binocular rivalry.

The mechanism by which respiration affects brain activity is not clear. However, the literature suggests two potential mechanisms for rhythmic coupling between the brain and internal organs. This occurs through direct sensory input, such as olfactory information transmitted by the olfactory bulb [10, 15], and through indirect neuromodulators influenced by internal organs, such as norepinephrine/noradrenaline (NE/NA) [5, 20, 23, 24, 27, 28, 34]. In this paper, we assume that the nonlinearity of ESN is modulated by NA relased by the LC which is further modulated by the respiratory activity [24, 34].

Hereinafter, we describe ESN dynamics, respiration dynamics, LC dynamics, respiration-LC coupling dynamics, and the LC-ESN coupling dynamics, in that order.

### ESN dynamics to describe binocular rivalry

The ESN dynamics associated with binocular rivalry is described as follows

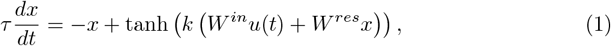

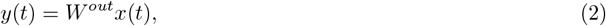

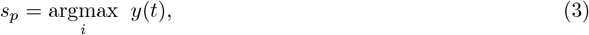

where *τ* is the time-constant of a neuron, *x ∈ R*^*N*^ describes the state vector of *N* neurons, *u ∈ R*^*L*^ is a input vector (*L* = 2) which describe the input received by the left and right eye and *u* = [1, 1]^*t*^. tanh(*z*) is the activation function of a neuron, *k* is modulated by NE [2, 30, 33], *W* ^*in*^ *∈ R*^*N×M*^ is the input matrix, *W* ^*res*^ *∈ R*^*N×N*^ is the weight matrix which depicts the connection between neurons, *W* ^*out*^ *∈ R*^*O×N*^ is the output matrix, and furthermore, *O* = 2. *s*_*p*_ is the index of the maximum output. Note that *s*_*p*_ is either 0 or 1 because *O* = 2, and we assumed *s*_*p*_ = 0 and 1 correspond to the perceptual state where the left-eye image is dominant, and right-eye image is dominant, respectively.

### Respiration dynamics

Respiration dynamics is generated from the integrated closed-loop system consisting of a mechanical lung, the gas exchange system of O_2_*/*CO_2_, a periodic heartbeat activity, and the central pattern generator to control the respiration-relevant muscles [11]. We changed the respiration frequency by adjusting the bias signal to the RTN nucleus.

### LC dynamics

LC neurons are modeled by Hodgkin-Huxley (HH) type neurons. The lateral neurons exert an inhibitory effect on a LC neuron, and furthermore, neighboring neurons are connected by a gap junction. More detail information about LC neuron are seen in our other manuscripts [11, 35].

### Respiration influence on LC dynamics

It is established that the activity of locus coeruleus (LC) neurons is subject to modulation by respiration [24]. Indeed, the LC receives direct synaptic projections from the preBötC [34]. Furthermore, LC neurons have chemical receptor for CO_2_*/*H^+^ [28], and receives synaptic connections from NTS which receives information of pulmonary stretch receptor [20]. Based on these evidences, we assumed that LC neurons receives respiratory-relevant information of pCO_2_, PSR, and pre-I/I neuron activity. The analog signals of pCO_2_, PSR, and pre-I/I neuron activity are input to the small neuron ensembles, each consisting of 5 Hodgkin-Huxley style neurons. In addition, the input gains of these respiratory signals are adjusted so that the HH ensembles have the same firing rate range.

The primary objective of this manuscript is to examine the impact of stationary respiration on binocular rivalry. Consequently, we exclude the influence of NE on breathing activity, despite the established role of LC in regulating breathing through the use of NE [5].

### *α*_+_ and *α*_*−*_ models of LC-ESN coupling via NA

LC neurons release NE in order to modulate the activation gain of ESN neurons. The effective amount of NE in ESN neurons is described as follows.

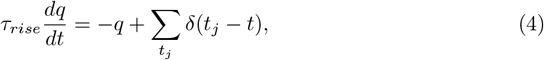

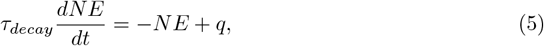

where *t*_*j*_ depicts the spike trains of LC neurons, *τ*_*rise*_ = 150 [ms] and *τ*_*decay*_ = 350 [ms]. The effect of norepinephrine (NE) on cortical dynamics is complex. In vitro observations indicate that NE can both increase and decrease the firing rate of neurons, depending on the specific brain region [3] and the volume of NE projected to specific neurons [1, 18]. A cortical neuron receives between 10^3^ and 10^4^ synaptic projections from other cortical neurons. Consequently, the global activity of cortical neurons is regarded as background noise to a neuron [4, 6], the intensity of which determines the nonlinearity of the neuron firing rate [7, 19]. Based on these considerations, two influence models of NE are proposed. In particular, we assumed that the parameter *k* in Eq. 2, which determines the nonlinearity of ESN neurons, is modulated by NE based on the following linear relationship:

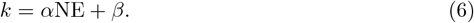

Note that the nonlinearity of ESN neurons increases with the value of the gain parameter, *k*. The positive value of *α, α >* 0, indicates that NE has the effect of reducing background noise and increasing the nonlinearity of a neuron, which is consistent with the model of Refs. [2, 30, 33]. Conversely, *α <* 0 assumes that NE enhances background noise activity of cortical neurons and increases the linearity of a neuron. We considered two parameter models; (*α, β*) = (0.07, *−*0.5) (*α*_+_ model); and (*α, β*) = (*−* 0.07, 3.65) (*α*_*−*_ model).

The cortical activity also exerts an influence of LC activity [2, 33]. Consequently, it was assumed that the mean activity of ESN is input to LC neurons., i.e., Σ tanh(*kx*) is input to 10 HH neuron ensemble and converted to spike-trains, the 10 HH neurons sent excitatory synaptic connections to LC neurons.

## 3 Results

We computed the likelihood function *L*(*ϕ*) of perceptual alternation with respect to the respiratory phase *ϕ* using *α*_+_ and *α*_*−*_ models. Fig. 2 and Fig. 3 show the likelihood function of the perceptual alternation with respect to the respiration phase *ϕ* and with different respiration frequencies (respiration count per minute). It is shown that the peak position of the likelihood function is weakly dependent on the respiration frequency. Furthermore, it is shown that the count of the perceptual alternation is dependent to the respiration frequency. This result is consistent to the previous study [37]. It is worth noting that the peak position of the likelihood distribution is around *−π/*4 and 3*/*4*π* regardless of the difference between *α*_+_ and *α*_*−*_.

**Figure 1.**
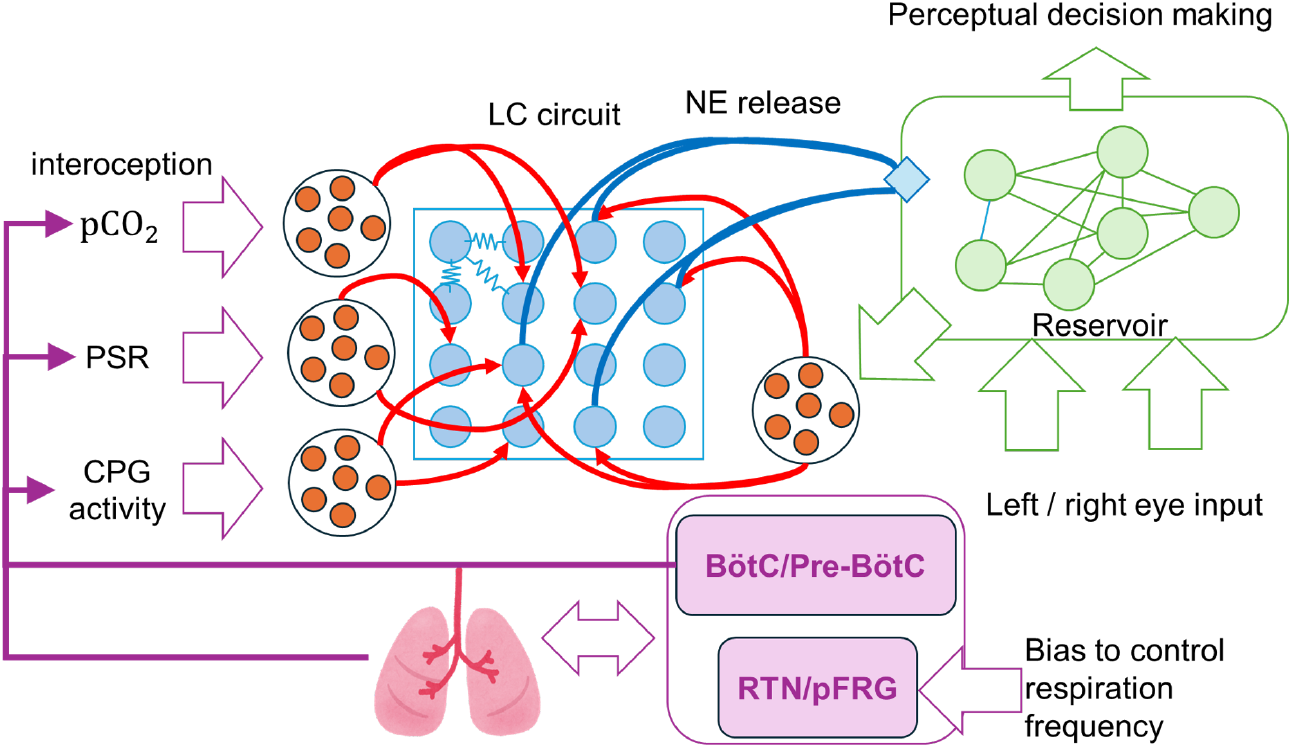
A coupling model of respiration, LC, and ESN, which is capable to reproduce the dependency of human binocular rivalry on respiration. Respiratory CPG (BötC/Pre-BötC and RTN/pFRG controls generate periodic movements of lung diaphragm. In the lung alveoli exchanges O_2_ and CO_2_. The CPG activity, the signal of pulmonary stretch receptor, and partial pressure of carbon dioxide pCO_2_ is passed to LC neurons. The neighbouring LC neurons are connected electrically by gap junctions, and inhibiting each other by the local emission of NE. LC neurons release NE to cortical reservoir neurons, and reservoir neurons send excitatory connections to LC neurons.

**Figure 2.**
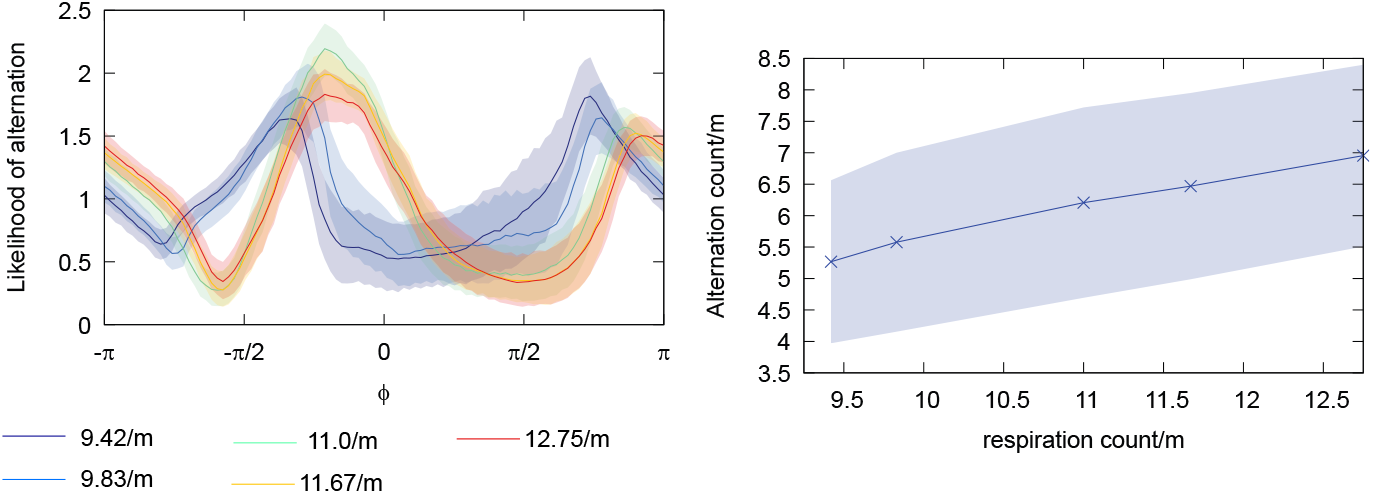
The likelihood function of perceptive alternation with respect to the respiration phase based on the *α*_+_ model. We collected 900 trials of 720 [s] numerical simulation. The color bars represent the standard deviations.

**Figure 3.**
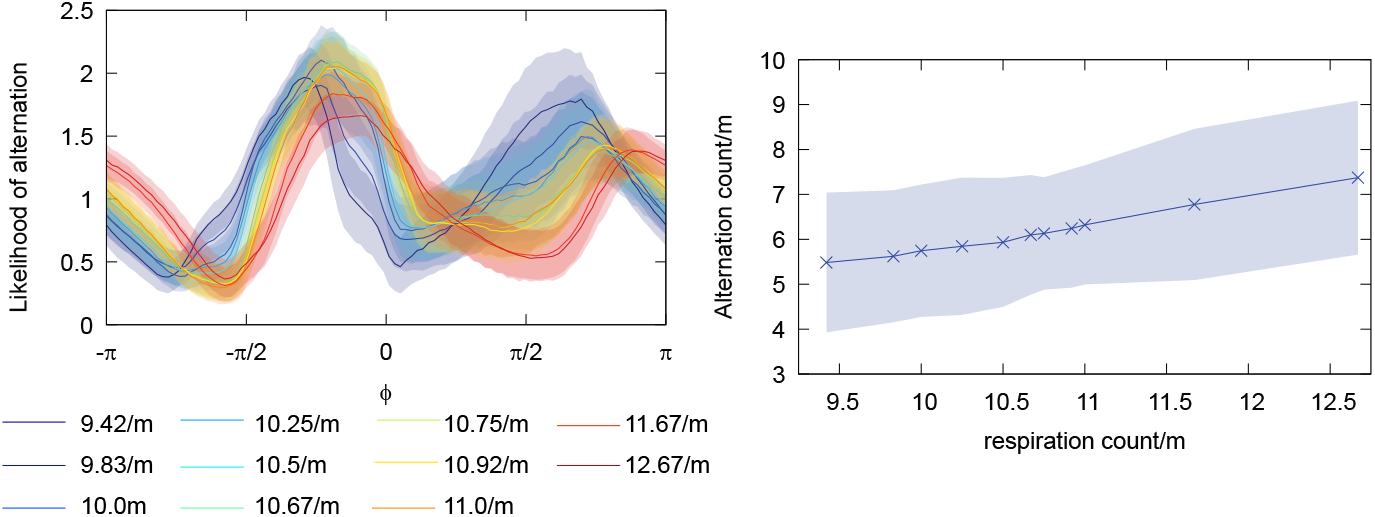
The likelihood function generated based of *α*_*−*_ model. It should be noted that there was no significant difference with *α*_+_ model. The color bars represent the standard deviations.

## 4 Conclusion

This paper describes a model of respiration-synchronized perceptual alternation in binocular rivalry. We found that reservoir dynamics can generate respiration-synchronized perceptual alternation when the reservoir system is coupled to respiratory dynamics via NA. The most important finding was that the likelihood peak phase is independent of whether the NA increases or decreases the nonlinearity of the neuron’s activation function. This suggests that the perceptual alternation is driven only by the respiratory phase information and not by the differential information of the respiratory signal. Further analysis of how respiration-related interoception induces perceptual alternation in binocular rivalry will be a future direction.

## Acknowledgements

This research is supported by Toyota Central R&D Labs., Inc.

